# Stathmin-2 regulates microtubule dynamics and axonal resilience in human motor neurons

**DOI:** 10.64898/2026.11.07.764673

**Authors:** Kaya J.E. Matson, Cecilia R. Petruconis, Amber Hernandez, Jay R. Dave, Suradip Das, Erika L.F. Holzbaur

**Author notes:** Corresponding author: Erika L.F. Holzbaur University of Pennsylvania Perelman School of Medicine 638A Clinical Research Building 415 Curie Boulevard Philadelphia, PA 19104.

## Abstract

Amyotrophic lateral sclerosis (ALS) is characterized by the selective degeneration of motor neurons, yet the mechanisms that either induce neurodegeneration or confer neuronal resilience remain poorly understood. Loss of the microtubule-associated protein Stathmin-2 (STMN2) is a downstream consequence of TDP-43 pathology present in nearly all cases of ALS and has emerged as a central feature of disease. However, the mechanisms by which STMN2 maintains homeostasis in motor neurons remain incompletely defined. Here, we investigated the role of STMN2 in regulating cytoskeletal organization and axonal function. Using TIRF microscopy at a single molecule resolution, we found that increasing STMN2 concentrations suppressed microtubule polymerization and increased catastrophe. In human induced pluripotent stem cell-derived motor neurons, STMN2 depletion increased baseline rates of microtubule polymerization. However, STMN2 depletion impaired recovery of axonal microtubules following transient nocodazole-induced depolymerization, indicating that STMN2 is required for efficient cytoskeletal remodeling after stress. Consistent with these findings, STMN2-depleted neurons exhibited impaired microtubule recovery following acute depolymerization and reduced axonal recoil after optically induced local microtubule severing. Finally, STMN2 depletion disrupted the formation of neuromuscular junction-like contacts in human motor neuron-myocyte co-cultures, linking impaired cytoskeletal homeostasis to loss of distal neuronal connectivity. Together, these findings identify STMN2 as a key regulator of cytoskeletal resilience that coordinates microtubule remodeling, mitochondrial transport, and maintenance of neuromuscular junctions. We propose that impaired resilience of the axonal cytoskeleton represents a critical point of vulnerability in motor neurons in patients with ALS, establishing cytoskeletal resilience as a potential therapeutic target downstream of TDP-43 pathology.

## INTRODUCTION

Amyotrophic lateral sclerosis (ALS) is a fatal neurodegenerative disease characterized by the selective degeneration of upper and lower motor neurons. More than 50 genes have been linked to ALS, with the majority of cases still classified as sporadic. Despite extensive effort, it remains unclear why motor neurons are particularly vulnerable. Their extraordinary morphology presents a fundamental challenge to neuronal maintenance as motor neurons extend axons that can exceed a meter in length, requiring sustained transport along the axon to maintain distal synaptic function.

Long-range axonal transport requires a dynamic microtubule cytoskeleton. Microtubules are polymers of tubulin heterodimers that serve as the primary tracks for kinesin- and dynein-driven cargo transport. While axonal microtubules were long considered to be relatively stable, live cell imaging reveals continual cycles of tubulin assembly and disassembly that support remodeling, particularly in response to physiological stress (*1*). Proper regulation of microtubule dynamics is therefore essential for maintaining axonal homeostasis, ensuring the appropriate distribution of organelles and macromolecules, and preserving neuromuscular junction (NMJ) integrity and function. Disruption of microtubule organization and axonal transport have been documented in both familial and sporadic ALS and are closely associated with axonal dysfunction (*2*). Understanding how motor neurons regulate axonal microtubule dynamics may reveal mechanisms that either preserve axonal integrity or make these cells selectively vulnerable to irreversible degeneration.

Pathological TDP-43 mislocalization, characterized by cytoplasmic accumulation and depletion from the nucleus, is observed in approximately 97% of ALS cases (*3*, *4*). Loss of nuclear TDP-43 function disrupts RNA processing in affected neurons, altering the expression and splicing of hundreds to thousands of transcripts (*4*, *5*). Among these, *Stathmin-2* (STMN2), which encodes a neuron-enriched tubulin-binding protein highly expressed in motor neurons (*6*), has emerged as a particularly well-characterized and functionally important TDP-43 target. Loss of nuclear TDP-43 permits inclusion of a cryptic exon in *STMN2* pre-mRNA, introducing a premature termination codon and polyadenylation signal that result in a truncated, nonfunctional transcript and reduced production of full-length STMN2 protein. Consequently, STMN2 depletion is a well-established downstream consequence of TDP-43 dysfunction in ALS, particularly in cases with TDP-43 pathology (*3*, *4*). STMN2 binds tubulin dimers and regulates microtubule dynamics, and its depletion has been linked to impaired axonal growth and regeneration, as well as motor dysfunction in experimental models (*7–9*). The therapeutic potential of restoring STMN2 expression is currently being evaluated in the ANQUR Phase 1/2 clinical trial of QurAlis’s STMN2-restoring splice-switching antisense oligonucleotide (ASO), QRL-201, in people with ALS.

STMN2 binds soluble tubulin through its conserved stathmin-like domain. Recent work has further shown that tubulin binding regulates STMN2 protein stability and membrane localization, suggesting that STMN2 function may be influenced by the balance between tubulin association and membrane binding (*8*). Studies of other stathmin-family proteins provide additional insight into the range of mechanisms through which these proteins can regulate microtubules. Biochemical analyses have shown that stathmin-family proteins can bind soluble α/β-tubulin heterodimers and inhibit tubulin polymerization, although individual family members differ in their tubulin-binding properties (*10*). The neuronal stathmin-family protein STMN2 inhibits microtubule assembly and can promote disassembly of preassembled microtubules in vitro; its overexpression also enhances nerve growth factor-induced neurite outgrowth in PC12 cells (*11*). Subsequent work demonstrated that STMN2 can bind directly to assembled microtubules and differentially regulate their plus- and minus-end dynamics, indicating that its effects cannot be explained solely by soluble tubulin sequestration (*12*). These studies underscore the diversity of microtubule-regulatory activities within the stathmin family, but do not establish that STMN2 shares all of these mechanisms. How STMN2-tubulin interactions contribute to microtubule remodeling and axonal maintenance remains unclear.

Motor neurons depend on microtubules to maintain axons that span extraordinary distances, yet these polymers must remain dynamic to support local remodeling and effective cargo transport. STMN2-dependent regulation of microtubules could therefore influence not only axonal architecture but also the ability of the axon to withstand cytoskeletal perturbation and maintain distal connectivity in response to cellular stress. We hypothesize that STMN2 regulates axonal microtubule dynamics by locally regulating the availability of soluble tubulin, thereby supporting localized cytoskeletal remodeling. This model predicts that STMN2 depletion induced by TDP-43 dysfunction may alter basal microtubule assembly while also compromising the ability of motor neurons to restore microtubule organization following cytoskeletal disruption. Because the microtubule cytoskeleton supports axonal trafficking, such changes could extend beyond cytoskeletal architecture to affect organelle localization and distal synaptic connectivity. In motor neurons, preserving these distal connections is particularly important for maintaining neuromuscular junctions (NMJs), specialized synapses that mediate communication between motor neurons and skeletal muscle.

Here, we investigated STMN2-dependent cytoskeletal regulation in human induced pluripotent stem cell-derived motor neurons (iMNs). Building on previous biochemical studies demonstrating that STMN2 binds tubulin and regulates microtubule dynamics, we found that STMN2 directly modulates microtubule polymerization in vitro, affecting both microtubule growth rates and catastrophe frequency. In iMNs, STMN2 depletion increased the ratio of polymerized to soluble tubulin, enhancing growth rates but impairing microtubule recovery following transient depolymerization. STMN2 depletion also diminished axonal recoil following light-induced localized microtubule severing with opto-katanin, revealing less robust axonal responses to cytoskeletal disruption. Paralleling these changes in cytoskeletal resilience, STMN2 depletion altered mitochondrial transport in the distal axon and impaired NMJ formation in motor neuron-muscle cocultures. Together, these findings identify STMN2 as a key regulator of microtubule dynamics in motor neurons that mediates the axonal response to cytoskeletal disruption. Our findings further link altered cytoskeletal regulation to changes in axonal transport and distal neuromuscular connectivity.

## RESULTS

### STMN2 binds tubulin and regulates microtubule dynamics in vitro

STMN2 belongs to the stathmin family of tubulin-binding proteins and contains a conserved tubulin-binding region (Figure 1A). To examine the molecular basis of STMN2-tubulin interactions, we generated an AlphaFold model of STMN2 bound to α/β-tubulin heterodimers. The model predicts that STMN2 engages two α/β-tubulin heterodimers, providing a structural framework for understanding how STMN2 may regulate tubulin availability (Figure 1B).

**Figure 1:**
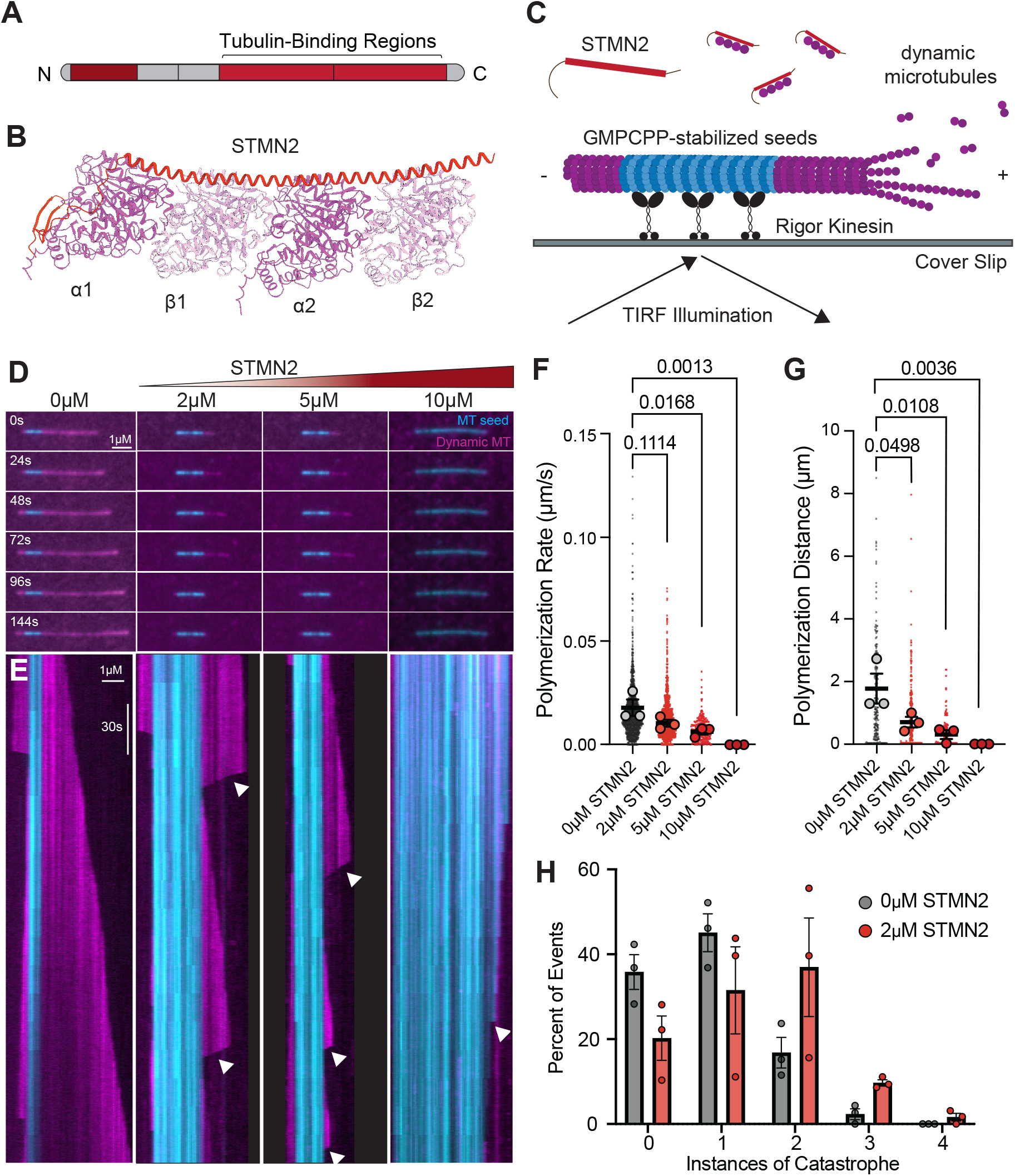
STMN2 binds soluble tubulin and regulates microtubule polymerization in vitro. (**A**) Domains of STMN2. (**B**) AlphaFold model of STMN2 bound to α/β-tubulin heterodimers. (**C**) Schematic of total internal reflection fluorescence (TIRF) microscopy, with GMPCPP-stabilized seeds (cyan) and free tubulin (magenta) in the presence or absence of recombinant STMN2 (red). (**D**) Time series of microtubule growth and catastrophe. Scale bar is 1µm. (**E**) Representative kymographs of microtubule growth and catastrophe over 3 minutes (3 frames/second). Instances of catastrophe indicated by white arrow. Scale bars are 1µm and 30 seconds. (**F**) Microtubule polymerization rate (µm/second). (**G**) Microtubule polymerization distance (µm). (**H**). Percent of events that resulted in 0, 1, 2, 3 or 4 events of catastrophe over the course of a 3-minute recording. Fisher’s exact test, *p* = 0.0013.

To interrogate the effects of STMN2 on microtubule dynamics, we used total internal reflection fluorescence (TIRF) microscopy to monitor microtubule growth from GMPCPP-stabilized microtubule seeds in the presence of 10µM soluble tubulin with increasing concentrations of recombinant STMN2 (Figure 1C, Supplemental Figure 1). Increasing concentrations of recombinant STMN2 reduced microtubule polymerization rates and growth distances (Figure 1D-G). At 2μM STMN2, catastrophe frequency increased, consistent with more frequent transitions from microtubule growth to shrinkage (*p* = 0.0013, Figure 1H). At higher STMN2 concentrations (10µM), microtubule growth was strongly suppressed with minimal detectable polymerization. We even noted shortening of GMPCPP-stabilized microtubule seeds at higher STMN2 concentrations (Figure 1E).

Previous biochemical studies of stathmin-family proteins have established that these proteins can inhibit microtubule assembly through interactions with soluble tubulin (*10*, *11*). However, work on STMN2 has also demonstrated direct regulation of assembled microtubules and distinct effects on plus- and minus-end dynamics (*12*). Thus, although our findings are consistent with STMN2-mediated regulation of soluble tubulin availability, they do not establish that tubulin sequestration is the sole mechanism through which STMN2 influences microtubule behavior.

Recent work has established that tubulin binding also regulates STMN2 protein stability and membrane localization (*8*), highlighting the potential for STMN2–tubulin interactions to coordinate cytoskeletal regulation with other cellular functions. Our findings extend this work by demonstrating that STMN2 directly modulates microtubule polymerization dynamics in a concentration-dependent manner.

These results demonstrate that STMN2 regulates microtubule dynamics in a concentration-dependent manner in vitro, consistent with a potent role for STMN2-mediated tubulin sequestration in shifting the balance between microtubule assembly and disassembly.

### STMN2 localizes throughout axons and is reduced in ALS

To investigate how STMN2 regulates microtubule dynamics in human motor neurons, we first examined its subcellular distribution to determine whether it is positioned to regulate distal axonal function. In iMNs, STMN2 RNA puncta detected using smFISH were concentrated in the soma and much less abundant in distal axons (Supplemental Figure 2A,B). Similarly, STMN2 protein was enriched in neuronal somas but was also detectable throughout axons, with pronounced enrichment at distal axon tips (Supplemental Figure 2C). To assess how STMN2 reaches these compartments, we tracked endogenously tagged STMN2 in human iPSC-derived excitatory neurons (Figure 2A). STMN2 moved bidirectionally along axons, with anterograde velocities consistent with translocation by the fast axonal motor KIF1A (Figure 2B-C). Together, these observations indicate that STMN2 protein is transported along axons to reach distal neuronal compartments, where it is positioned to regulate local cytoskeletal organization.

**Figure 2:**
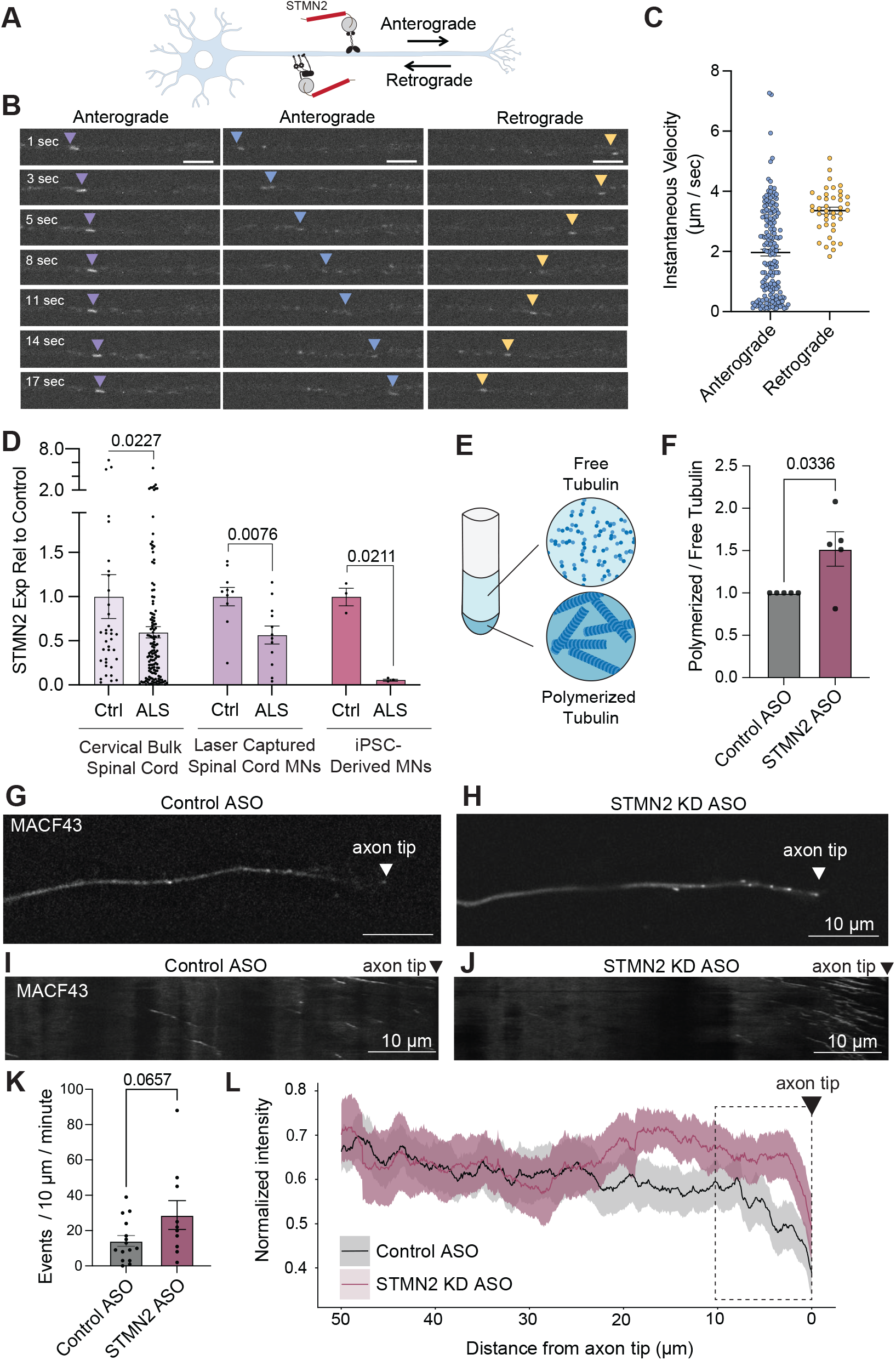
STMN2 regulates microtubule organization through soluble tubulin in human iPSC-derived motor neurons. (**A**) Cartoon showing the relative direction of anterograde and retrogradely transported STMN2 in iMNs. (**B**) Representative images showing STMN2 motility over time in iMN axons. Scale bar is 5µm. (**C**) Quantification of STMN2 motility, showing instantaneous velocity (µm/second) in the anterograde and retrograde direction in iMN axons. (**D**) STMN2 expression from published datasets including bulk spinal cord, laser captured motor neurons, and human iMNs. STMN2 expression levels in control samples were normalized to 1. (**E**) Cartoon depicting the ultracentrifugation and separation of free vs polymerized tubulin. (**F**) Quantified relative ratio of polymerized to free tubulin normalized to control ASO treatments. (**G-H**) Representative images of control ASO (**G**) and STMN2 ASO (**H**) axon tips transfected with GFP-MACF43. Comets represent growing ends of microtubules. Scale bar is 10 µm. (**I-J**) Representative kymographs of control ASO (**I**) and STMN2 ASO (**J**) MACF43 polymerization in axon tips. Scale bar is 10 µm. (**K**) MACF43 events per 10 µm per minute in the last 10 µm of the axon tip. P values represent values from unpaired t test. (**L**) An intensity summation over 3 minutes of MACF43 signal in the axon tip of DIV 21 iMNs. Control ASO average intensity is shown in black and STMN2 average intensity is shown in maroon. SEM is shown in 50% opacity lines above and below the average lines.

STMN2 depletion is a prominent consequence of the mis-splicing and resultant incorporation of cryptic exons induced by TDP-43 dysfunction and thus is predicted to be a common pathological hallmark across the majority of ALS cases. To further assess STMN2 expression across ALS contexts, we queried published transcriptomic datasets from human cervical spinal cord (*13*), laser-captured spinal motor neurons from sporadic ALS cases (*14*), and iMNs carrying a TARDBP (which encodes TDP-43) mutation (*15*). We noted significant reductions in STMN2 expression across these datasets (Figure 2D). This analysis further supports STMN2 depletion as a feature of ALS-associated motor neuron pathology.

### STMN2 depletion increases basal microtubule polymerization in human motor neurons

Since STMN2 suppressed microtubule growth in vitro, we hypothesized that reducing STMN2 would increase availability of soluble tubulin and promote microtubule assembly in neuronal axons. We tested this hypothesis by treating iMNs with antisense oligonucleotides (ASOs) targeting STMN2 (*16*). STMN2-targeting ASOs reduced STMN2 expression by 78% relative to control treatment (*p =* 0.0384, Supplemental Figure 2D). To examine the resulting effects on tubulin polymerization, we compared the distribution of tubulin between polymerized and soluble pools (*17*). STMN2 depletion significantly increased the ratio of polymerized to soluble tubulin (Figure 2E,F, Supplemental Figure 2E), consistent with a shift toward enhanced microtubule assembly.

We next examined microtubule organization at distal axon tips, where STMN2 is normally enriched. We noted that STMN2-depleted neurons exhibited broader axon tips (Supplemental Figure 2F). To measure microtubule growth dynamics, we performed live imaging with GFP-MACF43, a marker of growing microtubule plus ends (*18*). STMN2 depletion increased normalized MACF43 signal intensity in the distal 20μm of axon tips (Figure 2G-L). The number of detectable microtubule growth events in the most distal 10µm of the axon also showed an upward trend, although this difference did not reach statistical significance (*p* = 0.0657, Figure 2K). Thus, STMN2 depletion altered distal axon morphology and increased the local axon tip intensity MACF43 signal. Together with the increased polymerized-to-soluble tubulin ratio, these findings indicate that STMN2 depletion shifts basal microtubule organization toward polymerization.

### STMN2 depletion impairs microtubule recovery after cytoskeletal disruption

The increased rates and extent of microtubule growth in the axon tip observed under basal conditions prompted us to examine whether STMN2 depletion also affects microtubule recovery after acute disruption of microtubules. We transiently treated iMNs with nocodazole (100nM for 30min) to depolymerize microtubules and monitored polymerization following nocodazole washout (Figure 3A). In control axons, microtubule polymerization was halted during nocodazole treatment but rapidly recovered after nocodazole washout (Figure 3B-D,F). In contrast, STMN2-deficient neurons showed increased microtubule polymerization dynamics in the axon prior to nocodazole treatment, with markedly greater variability in growth rates observed (Figure 3D,F). Strikingly, following nocodazole-induced depolymerization and washout, STMN2-depleted neurons exhibited impaired polymerization recovery (Figure 2E,F). This impairment suggests that cytoskeletal recovery following transient stress is facilitated by the presence of tubulin-pools bound to STMN2 locally in the axon.

**Figure 3:**
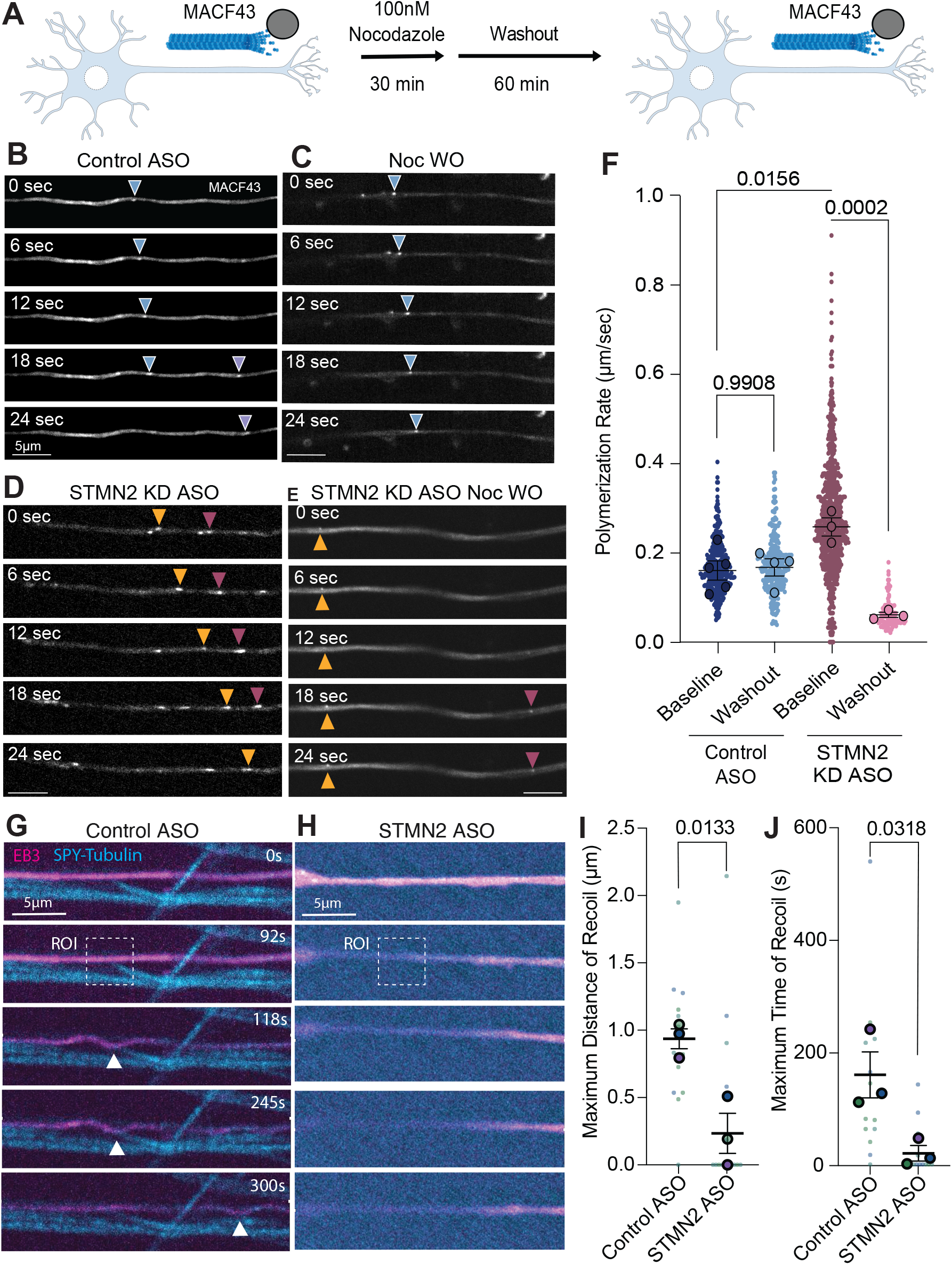
STMN2 depletion alters axonal responses to localized microtubule severing. (**A**) Cartoon showing experimental timeline of nocodazole treatment and washout of MACF43-positive DIV 21 iMNs. (**B-C**) Representative time series of MACF43 at baseline and following nocodazole washout in control ASO (**B,D**) and STMN2 ASO (**C,E**). (**F**) Polymerization rate of MACF43 at baseline and after nocodazole washout in control ASO- and STMN2 ASO-treated DIV 21 iMNs. P values represent one-way ANOVA with Šídák’s multiple comparisons test. (**G,H**) Representative time series of MACF43 in control ASO and STMN2 ASO treated iMNs transfected with EB3 (magenta) and optokatanin (not shown). SPY-tubulin dye (cyan) labels tubulin-dense axons. (**I**) Quantification of the maximum distance of recoil in µm, measured from original axon position at timepoint 0. (**J**) Quantification of the maximum time of recoil in seconds, measured from original axon position at timepoint 0. P values represent values from unpaired t test.

To determine whether these effects reflect a general consequence of altered STMN2 abundance, we also examined neurons overexpressing STMN2. Overexpression increased baseline microtubule polymerization rates (Supplementary Figure 3A-C). However, polymerization recovery after nocodazole washout did not differ from control conditions (Supplementary Figure 3C). Thus, both STMN2 depletion and overexpression altered baseline microtubule dynamics, whereas impaired recovery was observed following STMN2 depletion These findings distinguish the effects of STMN2 abundance on basal polymerization from its role in maintaining the capacity for cytoskeletal recovery after stress.

The impaired recovery of microtubule polymerization following nocodazole washout suggested that STMN2 depletion compromises the capacity of axonal microtubules to respond to cytoskeletal disruption. To further investigate this response using an independent perturbation, we used opto-katanin to induce localized microtubule severing through light-induced recruitment of the microtubule-severing enzyme katanin (*19*). We found that localized microtubule severing elicited a pronounced axonal recoil in control iMNs (Figure 3G,I,J). In contrast, STMN2-depleted iMNs exhibited reduced recoil distance and duration (Figure 3H,I,J). These findings suggest that STMN2 depletion alters the mechanical response of axons to acute microtubule disruption. The axonal deformation we observed following microtubule severing resembles the microtubule kinks associated with neurite retraction recently described by Lin et al (*20*) who showed that inhibiting ARP2/3-mediated actin branching induces microtubule buckling in developing neurons. Although these studies use distinct perturbations, both reveal a relationship between microtubule organization and neurite retraction. Together with the impaired recovery of microtubule polymerization following nocodazole washout, these findings indicate that STMN2 depletion alters axonal responses to acute cytoskeletal perturbation.

### STMN2 depletion alters trafficking and disrupts neuromuscular junction formation

Microtubules provide essential tracks for long-distance transport of organelles along the axon. We therefore asked whether STMN2 depletion alters mitochondrial movement in human motor neurons. We quantified mitochondrial motility in axons of wild-type motor neurons treated with control or STMN2-targeting ASOs. In control-treated neurons, most mitochondria (labeled with MitoTracker Deep Red) remained paused during imaging, consistent with stable mitochondrial docking observed in healthy axons. STMN2 depletion reduced the fraction of paused mitochondria and increased the fraction moving anterogradely, without a comparable increase in retrograde or bidirectional movement (Figure 4A-C). These findings indicate that STMN2 depletion alters both microtubule dynamics and organelle transport, raising the possibility that these changes can destabilize the distal axon.

**Figure 4:**
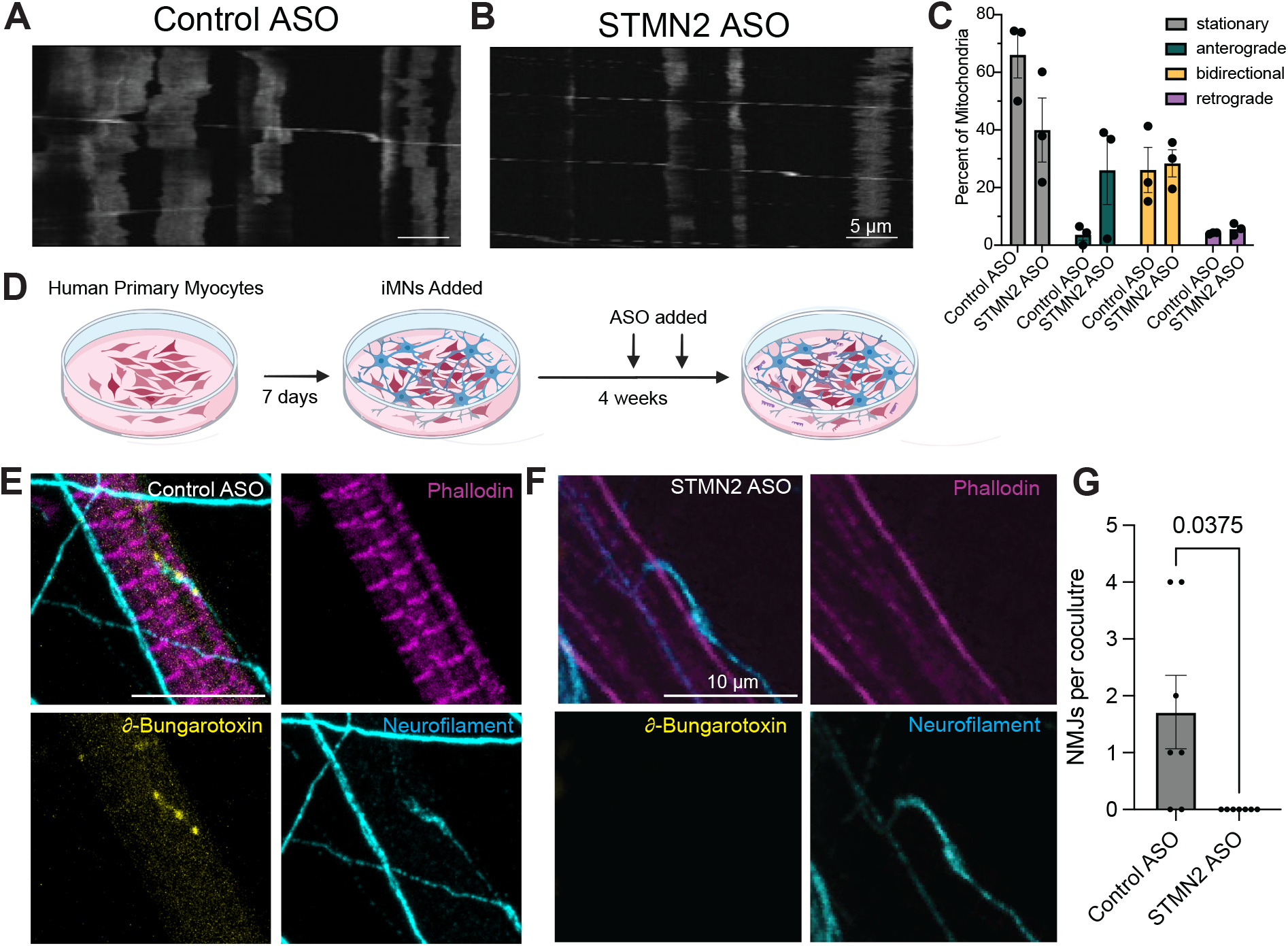
Loss of STMN2 disrupts neuromuscular junction formation. (**A-B**) Kymographs of with control ASO (**A**) and STMN2 ASO (**B**) treated mitotracker-red labeled DIV 21 iMNs. Scale bars are 10 µm. (**C**) Quantification of mitochondrial motility in control ASO- and STMN2-ASO-treated DIV 21 iMNs. (**D**) Cartoon graphic showing experimental setup of iPSC-derived myocytes and iMN co-cultures. (**E,F**) Immunostaining of fixed co-cultures with control ASO (**E**) and STMN2 ASO (**F**) treated conditions. Alpha-bungarotoxin is shown in yellow, phalloidin is shown in magenta and neurofilament is shown in cyan. Scale bars are 10 µm. (**G**) Quantification of the number of NMJs per co-culture. P values represent values from unpaired t test.

Specifically, we asked whether the altered cytoskeletal dynamics, mechanical response, and transport properties observed in STMN2-depleted axons correlate with impaired formation or stabilization of NMJs. To address this question, human iPSC-derived motor neurons were co-cultured with human primary skeletal myocytes to generate functional NMJs (*21*). Myocytes were differentiated for seven days prior to addition of motor neuron neurospheres (*22*), and co-cultures were maintained for four weeks to allow NMJ maturation (Figure 4D). Control or STMN2-targeting ASOs were administered during weeks two and three of the co-culture period.

Following maturation, cultures were fixed and stained with α-bungarotoxin to label postsynaptic acetylcholine receptor clusters together with neurofilament and phalloidin to visualize presynaptic motor axons (Figure 4E,F). While control cultures formed α-bungarotoxin-positive neuromuscular contacts, STMN2-depleted cultures did not form α-bungarotoxin-positive contact sites (*p* = 0.0375, Figure 4G). These findings indicate that STMN2 depletion disrupts the formation or maintenance of neuromuscular contacts in human motor neuron-myocyte cocultures, extending the consequences of STMN2 loss beyond axonal cytoskeletal organization to distal neuromuscular connectivity.

## DISCUSSION

Motor neurons must maintain an extensive axonal cytoskeleton while retaining the capacity for efficient remodeling in response to cellular stress over a lifetime. How these neurons preserve the capacity for cytoskeletal remodeling throughout the lengths of their exceptionally long axons remains unclear. Here, we identify STMN2 as a critical regulator of microtubule dynamics and axonal homeostasis in human motor neurons. STMN2 depletion increased basal microtubule polymerization but impaired recovery following cytoskeletal disruption, revealing a distinction between steady-state microtubule assembly and the capacity to remodel after stress. These changes induced by STMN2 depletion were accompanied by altered mitochondrial transport and impaired NMJ formation. Together, our findings suggest that STMN2 supports axonal resilience by maintaining the capacity for appropriate cytoskeletal remodeling, rather than simply controlling the steady-state level of microtubule assembly.

STMN2 has emerged as a critical downstream effector of TDP-43 dysfunction in ALS. TDP-43 pathology, characterized by nuclear clearance and cytoplasmic accumulation, is observed in approximately 97% of ALS cases, spanning most sporadic and familial disease. Loss of nuclear TDP-43 induces cryptic splicing and premature polyadenylation of STMN2 transcripts, reducing functional STMN2 expression in human motor neurons (*4*, *16*). Consistent with published studies, our analysis of transcriptomic datasets revealed reduced STMN2 expression in human spinal cord, laser-captured motor neurons from sporadic ALS cases, and TARDBP-mutant iMNs (Figure 2A), supporting the relevance of STMN2 depletion across distinct ALS contexts. Previous work by Klim et al. and Baughn et al. established the functional importance of this pathway by demonstrating that restoring STMN2 expression can rescue aspects of axonal regeneration and lysosomal trafficking in TDP-43-deficient human motor neurons (*4*, *16*). These studies provided compelling evidence that STMN2 loss is not merely a marker of TDP-43 dysfunction but a functionally important contributor to neuronal pathology. However, although this work established a role for STMN2 in neuronal regeneration and intracellular trafficking, it did not fully resolve how STMN2 supports the ongoing maintenance of the axonal cytoskeleton. Our findings extend this framework by identifying altered microtubule remodeling, mitochondrial transport, and distal synaptic connectivity as additional processes associated with STMN2 depletion. This broader functional perspective is particularly relevant to ALS, where the consequences of reduced STMN2 must be understood in the context of sustained axonal maintenance.

A notable observation from this study is the distinction between basal microtubule dynamics and recovery following cytoskeletal stress. Although STMN2 depletion modestly increased microtubule polymerization under steady-state conditions, these neurons exhibited a marked defect in rebuilding the microtubule cytoskeleton following transient nocodazole-induced depolymerization. Conversely, STMN2 overexpression also increased baseline polymerization but did not impair recovery after stress. Thus, the relationship between STMN2 abundance and basal polymerization is distinct from its contribution to microtubule recovery following perturbation. These findings indicate that increased polymerization alone does not necessarily reflect improved cytoskeletal function. Rather, STMN2 appears to support the capacity of motor neurons to dynamically remodel their microtubule network following disruption.

One possible explanation is that STMN2 binding to soluble tubulin influences the availability of tubulin for microtubule assembly. In this model, STMN2 may buffer the soluble tubulin pool, influencing basal polymerization while helping maintain the capacity for microtubule reassembly following perturbation. Our TIRF experiments support a direct role for STMN2 in regulating microtubule polymerization dynamics. However, our work does not establish that altered soluble tubulin availability is the sole mechanism underlying these effects.

Previous biochemical studies established that stathmin-family proteins bind soluble tubulin dimers and inhibit polymerization, providing a biochemical basis for tubulin buffering (*10*). Studies of STMN2, historically referred to as SCG10, further demonstrated that this neuronal stathmin-family protein can regulate microtubule dynamics through interactions with assembled microtubules. In particular, Manna et al. found that STMN2 promotes plus-end growth while increasing minus-end shortening and catastrophe, with these effects persisting under steady-state conditions in which soluble tubulin concentration was maintained (*12*). These findings demonstrate that individual microtubule dynamic parameters can be regulated independently and that the effects of a stathmin-family protein cannot necessarily be explained by changes in the soluble tubulin pool alone.

These studies also highlight the importance of distinguishing among stathmin-family members. Despite substantial sequence similarity, STMN2 and STMN1 differ in their effects on microtubule-end dynamics (*10*) and their ability to sequester soluble tubulin (*12*). Earlier work showed that STMN2 can inhibit microtubule assembly and promote disassembly of preassembled microtubules, while enhancing NGF-induced neurite outgrowth in PC12 cells (*11*). Together, these studies illustrate that microtubule destabilization is not inherently detrimental; regulated assembly and disassembly can support neuronal remodeling. However, the distinct effects of individual stathmin-family members on microtubule dynamics underscore the importance of defining STMN2-specific mechanisms rather than extrapolating from studies of other family members. Our findings extend this literature by identifying a role for STMN2 in maintaining microtubule recovery following transient depolymerization in human motor neurons. Although the molecular interactions underlying this effect remain to be determined, these results suggest that STMN2 contributes to cytoskeletal resilience by supporting dynamic remodeling rather than simply maintaining steady-state microtubule stability.

Previous work by Beccari et al. (2025) provided important evidence that STMN2 promotes motor axon regeneration and supports neuromuscular reinnervation and functional recovery following injury (*23*). Notably, their demonstration that a tubulin-binding-deficient STMN2 variant could rescue axon regeneration challenged the prevailing model that STMN2’s regenerative function is mediated exclusively through its interaction with tubulin. However, this finding does not preclude a distinct role for STMN2-tubulin interactions in other aspects of neuronal homeostasis, including the ongoing maintenance and remodeling of the axonal cytoskeleton. In parallel, Nolan et al. (2026) reported that pharmacological and genetic inhibition of the mevalonate pathway activates an ATF3-STMN2 regenerative program (*24*), further highlighting the integration of STMN2 into upstream signaling pathways that regulate neuronal regenerative capacity. Together, these studies position STMN2 within a broader regulatory network linking cellular stress responses, regeneration, and neuronal maintenance. Our findings extend this framework by identifying altered microtubule remodeling, mitochondrial transport, and distal synaptic connectivity as additional processes associated with STMN2 depletion. This broader functional perspective is particularly relevant to ALS, where the consequences of reduced STMN2 must be understood in the context of sustained axonal maintenance.

Recent work in from Lin et al. (2026) provides a complementary perspective on the relationship between microtubule organization and neuronal morphology (*20*). In developing neurons, inhibition of ARP2/3-mediated actin branching induced microtubule kinks associated with neurite retraction, suggesting that actin branching normally counteracts contractile forces that compress and buckle microtubules. Our opto-katanin experiments extend this context to human motor neurons by showing that localized microtubule severing elicits axonal recoil and that this response is altered by STMN2 depletion. Whereas Lin et al. manipulated actin branching to investigate the mechanical constraints on microtubules, our experiments directly perturb the microtubule network. The reduced recoil distance and duration following STMN2 depletion suggest that STMN2 influences the axonal response to acute microtubule disruption, although the underlying mechanism remains unresolved. Together with the impaired recovery of microtubule polymerization following nocodazole washout, these findings support a role for STMN2 in maintaining axonal cytoskeletal homeostasis under stress.

Recent work by Deng et al. (2025) provides a complementary framework for understanding how STMN2-tubulin interactions may contribute to this function (*8*). They showed that tubulin binding stabilizes STMN2 and reduces its association with trans-Golgi network membranes, leading them to propose that STMN2 transitions between soluble, tubulin-bound and membrane-bound states. They further suggested that spatial differences in soluble tubulin availability could influence STMN2 localization and function. Our findings extend this framework by demonstrating that STMN2 regulates microtubule polymerization dynamics and is required for efficient recovery following cytoskeletal disruption in human motor neurons. Together, these studies raise the possibility that STMN2–tubulin interactions coordinate cytoskeletal regulation with the capacity to maintain axonal integrity under stress. Whether compartment-specific changes in STMN2–tubulin binding directly contribute to these responses remains to be determined.

The alterations in microtubule organization induced by STMN2 depletion that we observed were accompanied by changes in mitochondrial transport; specifically, a reduction in the fraction of paused mitochondria and an increase in anterograde movement, suggesting a shift in the balance between mitochondrial anchoring and active transport. In healthy motor neurons, mitochondria are positioned at sites of high metabolic demand, where they support ATP production and calcium buffering. STMN2-dependent changes in the microtubule cytoskeleton may therefore influence mitochondrial positioning by altering the balance between pausing and transport. Consistent with the potential functional consequences of altered mitochondrial positioning, a study of human induced pluripotent stem cell-derived neurons from individuals with bipolar disorder found increased axonal mitochondrial motility alongside reduced presynaptic mitochondrial retention and impaired activity-dependent ATP maintenance and calcium buffering (*25*). Although this study examined a different disease context and neuronal population, it illustrates that increased mitochondrial motility does not necessarily indicate improved cargo delivery or synaptic support. Whether STMN2 depletion compromises mitochondrial retention or bioenergetic function at distal motor axons and NMJs remains to be determined.

STMN2 depletion also impaired NMJ formation in human motor neuron-myocyte cocultures, linking altered cytoskeletal homeostasis to distal synaptic connectivity. This finding is consistent with the established importance of STMN2 in axonal growth and maintenance and with evidence that STMN2 loss can contribute to motor and sensory denervation (*26*). The NMJ is a particularly demanding site for axonal maintenance because its assembly and function require coordinated delivery of synaptic components, organelles, and cytoskeletal machinery to distal motor axons. Defects in microtubule remodeling or cargo positioning could therefore compromise NMJ formation or maintenance.

STMN2 abundance is also dynamically regulated by cellular stress, providing a potential link between stress responses and cytoskeletal homeostasis. Previous work demonstrated that stress-activated JNK signaling promotes STMN2 phosphorylation and degradation, suggesting that STMN2 turnover may contribute to the acute regulation of microtubule dynamics in injured neurons (*27*). More recently, Ellis et al. (2026) showed that acute cellular stress induces STMN2 depletion through proteasomal degradation, phosphorylation, and stress granule-mediated translational repression, whereas chronic translation deficits can sustain STMN2 loss independently of TDP-43-mediated splicing dysfunction (*28*). Although the relative contribution of these mechanisms to STMN2 depletion in ALS remains unclear, these findings suggest that STMN2 abundance may be influenced by both RNA processing and protein homeostasis. Our findings demonstrate that reduced STMN2 abundance compromises the capacity of human motor neurons to remodel their axonal cytoskeleton following stress, suggesting a potential link between TDP-43-associated STMN2 loss and impaired axonal resilience.

An additional consideration is that STMN2 abundance may be regulated at multiple levels during neuronal stress. Although TDP-43 dysfunction is a major mechanism of STMN2 transcript loss in ALS, recent work found that neuronal stress can also reduce STMN2 protein through translational repression, stress granule-associated mechanisms, and proteasomal degradation, independently of TDP-43-dependent RNA processing (*28*). Furthermore, reduced STMN2 expression sensitizes neuronal cells to stress, whereas moderately increasing STMN2 can be protective. More broadly, these results support a conceptual shift in how STMN2 function is viewed. Rather than acting solely as a regulator of basal microtubule dynamics, our data indicate that STMN2 promotes the ability of motor neurons to restore cytoskeletal organization following stress while maintaining balanced intracellular transport and distal neuronal function. This framework emphasizes cytoskeletal resilience as a dynamic cellular property that extends beyond maintenance of steady-state microtubule architecture.

In the context of ALS, where STMN2 expression is disrupted downstream of TDP-43 pathology in most patients, impaired cytoskeletal resilience may represent a convergent mechanism contributing to motor neuron vulnerability. Failure to efficiently restore microtubule organization following cytoskeletal stress could compromise axonal transport, disrupt mitochondrial positioning, and ultimately impair NMJ integrity. These findings highlight the importance of understanding how STMN2 supports long-term axonal maintenance and suggest that restoring STMN2 expression could preserve neuronal function beyond promoting axonal regeneration. Encouragingly, this therapeutic strategy is now being evaluated clinically in the ANQUR Phase 1/2 trial of QRL-201, an investigational antisense oligonucleotide designed to restore STMN2 expression by correcting aberrant RNA processing. Although interim results provide encouraging evidence of target engagement, the clinical benefit of STMN2 restoration remains to be established in adequately powered efficacy trials. By identifying cytoskeletal resilience as an important function of STMN2, our findings provide a mechanistic framework for investigating whether therapeutic restoration of STMN2 can preserve axonal integrity and distal connectivity in people living with ALS.

## Materials and Methods

### Human iPSC-derived motor neuron culture and differentiation

Human induced pluripotent stem cell (iPSC)-derived motor neurons (iMNs) were generated using as described previously (*29–31*). In short, iPSCs were maintained on Matrigel-coated culture dishes in mTeSR1 medium. For neuronal induction, iPSCs were dissociated using Accutase and plated in induction medium consisting of DMEM/F-12 supplemented with N2, non-essential amino acids, and GlutaMAX, containing 2 μg/mL doxycycline and 10 μM ROCK inhibitor Y-27632. Their media was changed daily.

On day 3, pre-differentiated iMNs were harvested using Accutase and cryopreserved in iMN freezing medium. For neuronal culture, 35-mm glass-bottom imaging dishes were coated with poly-L-ornithine. On day 4, cryopreserved pre-differentiated iMNs were thawed, washed, and plated onto coated imaging dishes in Neurobasal medium (Gibco; 21103049), 1% N2, 1% NEAA, 1% Optimal-Culture-One supplement (Gibco; A3320201), with 1× B27 (Thermo Fisher Scientific, Cat. #17504-044), BDNF (10 ng/mL), GDNF (10 ng/mL), NT-3 (10 ng/mL), and laminin (1 μg/mL), Following plating, cultures were maintained at 37°C in a humidified incubator with 5% CO₂. The following day, the medium was replaced with fresh iMN culture medium, and half medium changes were performed three times a week.

Neurospheres were generated using a modified protocol based on Whittaker et al. described (*32*). Briefly, iPSCs containing a doxycycline-inducible NIL cassette were dissociated and plated at a density of 7,000-10,000 cells per well in 384-well plates in neuronal induction medium supplemented with doxycycline. Cells were maintained under neuronal induction conditions for 4 days, with 50% medium changes performed daily. On day 4, the resulting neurospheres were gently collected by pipetting and transferred to dishes containing motor neuron maintenance medium for gentle washing and inspection. Neurospheres were subsequently transferred to either poly-L-ornithine (PLO)-coated imaging dishes for monoculture experiments or imaging dishes containing primary myocytes for co-culture experiments.

Human iPSCs with endogenously tagged STMN2-mScarlet and additional safe-harbor integration of doxycycline-inducible NGN2 were a gift from M. Ward (Berkeley, CA, USA). These iPSCs were maintained on Matrigel-coated culture dishes in mTeSR1 medium and differentiated into iNeurons as previously described (*29*, *30*). In short, iPSCs were maintained on Matrigel-coated culture dishes in mTeSR1 medium. iPSCs were dissociated using Accutase and plated in induction medium consisting of DMEM/F-12 supplemented with N2, non-essential amino acids, and GlutaMAX, containing 2 μg/mL doxycycline and 10 μM ROCK inhibitor Y-27632. On days 1 and 2 following induction, cells were fed with induction medium containing 2 μg/mL doxycycline without ROCK inhibitor. On day 3, pre-differentiated iNeurons were harvested using Accutase and cryopreserved in iNeuron freezing medium.

For neuronal culture, 35-mm glass-bottom imaging dishes were coated with poly-L-ornithine. Cryopreserved pre-differentiated iNeurons were thawed, washed, and plated in iNeuron culture medium consisting of BrainPhys Neuronal Medium (STEMCELL Technologies, Cat. #05790) supplemented with 1× B27 (Thermo Fisher Scientific, Cat. #17504-044), BDNF (10 ng/mL), NT-3 (10 ng/mL), and laminin (1 μg/mL), as described previously (30). Cultures were maintained at 37°C in a humidified incubator with 5% CO₂. The following day, the medium was replaced with fresh iNeuron culture medium, and half-medium changes were performed twice weekly. Neurons were maintained for 21 days after plating before live-cell imaging experiments.

### Antisense oligonucleotide-mediated STMN2 depletion

To reduce STMN2 expression, iMNs were treated with an STMN2-targeting antisense oligonucleotide (ASO; GGTCTTAGTCAAGCTCAGAG) or a non-targeting control ASO (CCTATAGGACTATCCAGGAA). Unless otherwise indicated, ASOs were applied at a final concentration of 5 µM for 7 days before analysis. For microtubule plus-end imaging using GFP-MACF43, ASOs were added 4 days before imaging. ASO sequences and chemical modifications were designed as described previously (*16*).

### RNA isolation and quantitative PCR

Total RNA was isolated from approximately 1 million iMNs using TRIzol (ThermoFisher, Cat# 15596026) to extract RNA according to the manufacturer’s instructions. Quantitative PCR was performed using Brilliant III Ultra-Fast SYBR Green qRT-PCR Master Mix (Cat# 600886) with primers targeting human STMN2 and GAPDH. Relative gene expression was calculated using the ΔΔCt method and normalized to GAPDH.

### Plasmid transfection

iMNs were transfected using Lipofectamine Stem (Thermo Fisher Scientific) and Opti-MEM (Gibco), following the manufacturer’s instructions and previously published neuronal transfection protocols. Briefly, plasmid DNA was combined with Lipofectamine Stem (STEM00001; Thermo Fischer Scientific) in Opti-MEM and incubated for 15 minutes before addition to neuronal cultures. For co-transfection experiments, the total amount of DNA was 0.4-1.5 µg per dish, with individual plasmids included at the ratios specified for the relevant experiment. Transfection reagents were removed after 1.5 hours, and neurons were imaged 48-72 hours later. For optogenetic katanin experiments, neurons were co-transfected with 0.8 μg EB3N-VVD-mCl3-iLID, 1.5 μg SspB-mCh-p60. Microtubule plus-end dynamics were assessed by transfecting GFP-MACF43 or EB3-mCherry in iMNs at 1µg per dish. Neurons were transfected with GFP-MACF43 and treated with control or STMN2-targeting ASO for 4 days before imaging. A detailed protocol for culturing and transfecting iPSC-derived neurons for live imaging is available on Protocols.io (*33*).

### Optogenetic recruitment of katanin

Optogenetic recruitment of katanin was performed as previously described (*19*). Localized activation was performed using 488-nm illumination pulses of 113 ms duration delivered every 4 seconds. These pulse settings were used as closely as possible to the published conditions.

### Live-cell fluorescence microscopy

Live-cell imaging was performed using an inverted Nikon Eclipse Ti microscope equipped with an UltraView Vox spinning-disk confocal system and an environmental chamber maintained at 37°C. Images were acquired using a 100x oil-immersion objective and a Hamamatsu ORCA-Fusion camera controlled by VisiView software. Imaging medium consisted of Hibernate A supplemented with 2% B27, BDNF (10 ng/mL), GDNF (10 ng/mL), NT-3 (10 ng/mL), and laminin (1 ng/mL). For axonal imaging, individual neurons were selected based on neuronal morphology and the presence of an identifiable axon, at least 200µm from the soma.

### Nocodazole-induced microtubule depolymerization and recovery

To assess microtubule recovery following pharmacological depolymerization, iMNs were treated with 100 nM nocodazole for 30 min at 37°C (*34*). Nocodazole was subsequently removed by three washes with motor neuron media, and neurons were returned to fresh motor neuron media for recovery. Live-cell time-lapse images were acquired at 1 frame per second for 5 minutes.

### Mitochondrial motility imaging and analysis

Mitochondrial motility was assessed by treating live iMNs with 1:10,000 dilution MitoTracker Deep Red FM (Thermo Fisher Scientific, M22426). Live-cell time-lapse images were acquired at 1 frame per second for 5 minutes. Images and movies were acquired within 1 hour of MitoTracker addition to minimize toxicity.

Kymographs of mitochondrial motility were generated by tracing a segmented line and using Multi Kymograph (width 3) in ImageJ. Mitochondrial movement was classified as paused, anterograde, or retrograde as previously described (*35*). In short, motile mitochondria were scored as anterograde (net displacement >10 μm in the anterograde direction within the 5-minute time lapse duration), retrograde (net displacement >10 μm in the retrograde direction), or bi-directional (net displacement <10 μm in either direction but total displacement >10 μm). Mitochondria were scored as stationary if net and total displacement were <10 μm. A pause was defined as a single or consecutive instantaneous velocity value of <0.083 μm/s.

### Tubulin polymerization and soluble tubulin fractionation

The relative distribution of polymerized and soluble tubulin was assessed using the Microtubule/Tubulin In Vivo Assay Biochem Kit (Cytoskeleton Inc) following manufacturer’s instructions. In short, approximately 1 million iMNs were collected and homogenized in lysis and microtubule stabilization buffer. Samples were centrifuged at 1,000 g for 5 minutes at 37°C. The supernatant was centrifuged at 100,000 g for 60 minutes at 37°C to separate soluble tubulin in the supernatant from polymerized tubulin in the pellet. The soluble fraction was collected, and the pellet fraction was resuspended in microtubule destabilization buffer. Tubulin was detected on a western blot using alpha/beta tubulin (Cat# 2148S, Cell Signaling Technology). The relative abundance of polymerized and soluble tubulin was assessed by calculating the ratio of polymerized to soluble tubulin.

### In vitro microtubule dynamics by TIRF microscopy

In vitro microtubule dynamics were assessed using purified human recombinant STMN2 protein (OriGene, TP302914; NM_007029) and dual-color microtubules by total internal reflection fluorescence (TIRF) microscopy. Tubulin was obtained from (Porcine Brain, Cat# T240-DX, Cytoskeleton Inc). GMPCPP-stabilized microtubule seeds were prepared by combining unlabeled tubulin and HiLyte 488-labeled tubulin (Cytoskeleton) to a final concentration of 50 μM. This mix was clarified at 352,000 × *g* for 10 min at 4 °C and kept on ice.

Flow chambers were prepared using silanized #1.5 glass coverslips (Warner) attached to glass slides (FisherScientific) using vacuum grease. To reduce non-specific binding, coverslips were cleaned through rounds of sonication in acetone, potassium hydroxide, and ethanol, dried, plasma cleaned and silanized with PlusOne Repel-Silane (GE Healthcare). Flow chambers were coated with 0.5 µM rigor kinesin-1E236A, washed and blocked with 5–8% Pluronic F-127 (Sigma). After washing, 25 nM TRITC Rhodamine-labeled GMPCPP-stabilized microtubules were flowed-in and left to attach to the coverslip for 1 min. Unbound microtubules were washed out with P12 (12 mM PIPES, 1 mM EGTA, 2 mM MgCl2). A final solution with STMN2 in BRB80, 10 μM tubulin mix, 1 mM Mg-GTP, 1 mM Mg-ATP in Dynamic Assay Buffer (P12, 0.3 mg/mL BSA, 0.3 mg/mL casein, 10 mM DTT, 15 mg/mL glucose, 0.05% methylcellulose) and an oxygen scavenging system (0.5 mg/mL glucose oxidase, 470 U/mL catalase; Sigma) was then flowed-in and let to equilibrate for ∼5 min at 37 °C before time lapse acquisition was initialized.

Dual-color time-lapse imaging was performed using a Nikon Eclipse Ti inverted microscope equipped with an UltraView Vox TIRF system and a 100×, 1.49 NA oil-immersion objective. Dynamic microtubule images were acquired at 3 frames per second, with microtubule seed images acquired every 13.3 seconds (once every 40 frames) for 3 min at 37°C. At least five videos were acquired for each chamber.

Kymographs of the dynamic microtubule and microtubule seed channels were generated for each microtubule at its maximum length by tracing a segmented line and using Multi Kymograph (width 3) in ImageJ. All microtubules where the entire length of the microtubule were in the field of view were analyzed. The microtubule plus-end was identified as the end with higher rates of growth and catastrophe in the 3-minute video. Each microtubule growth event was tracked using a segmented line to determine polymerization rate and distance.

### Neuromuscular junction co-culture

To assess the effects of STMN2 depletion on motor neuron–muscle interactions, human myocytes were differentiated and co-cultured with iMN neurospheres. Myocytes were cultured for 7 days before addition of iMN neurospheres. Co-cultures were maintained for an additional 4 weeks before fixation and immunostaining. Neuromuscular junction-like contacts were visualized by labeling neuronal processes with neurofilament, postsynaptic acetylcholine receptors with fluorescent α-bungarotoxin, and muscle cells with phalloidin.

### Immunofluorescence staining

For immunofluorescence analysis, cultured neurons were fixed using 4% PFA for 10 minutes and washed with phosphate-buffered saline (PBS). Cells were permeabilized with 0.1% triton for 10 minutes then blocked for an hour in 5% goat serum and 1% BSA in PBS. Cells were incubated overnight at 4°C with the respective primary antibodies in blocking solution. Next, cells were washed with PBS three times before incubation with respective secondary antibodies in blocking solution for 1 hour at room temperature. After 3 washes with PBS, samples were mounted in ProLong Gold Antifade Mountant (Life Technologies, P36930). Images were acquired as z stacks at 200 nm step-size using a PerkinElmer UltraView Vox spinning disk confocal on a Nikon Eclipse Ti Microscope. Fixed cell experiments were performed on an Apochromat 100x 1.49NA oil-immersion objective.

### Statistical analysis

Statistical analyses were performed using GraphPad Prism and R. Data are presented as mean ± SEM. The number of independent biological replicates and technical replicates is indicated in the corresponding figure legends. Unless otherwise indicated, statistical analyses were performed using independent biological replicate means as the unit of analysis.

### Author Contributions

K.J.E.M. and E.L.F.H. conceived and designed the study and wrote the manuscript. K.J.E.M., A.H., C.R.P., and J.R.D. performed experiments and analyzed data. S.D. and E.L.F.H. supervised the methodology and project administration.

## Acknowledgements

We thank Mariko Tokito for technical assistance and members of the Holzbaur lab for their valuable feedback. We thank Nicholas Palmer and Roberto Dominguez for help with AlphaFold modeling of STMN2. We thank Michael Ward and James Hawrot for the STMN2-mScarlet iPSC line. This work was supported by the ALS Finding a Cure, the Healey Center and FightMND via the ALS Scholars in Therapeutics Fellowship to K.J.E.M. and National Institutes of Health Grants NIGMS R35 GM126950 to E.L.F.H.

