## SupplementalFigures for "Stathmin-2 regulates microtubule dynamics and axonal resilience in human motor neurons"

**Supplemental Figure 1. Dose-dependent impact of STMN2 on microtubule polymerization.**

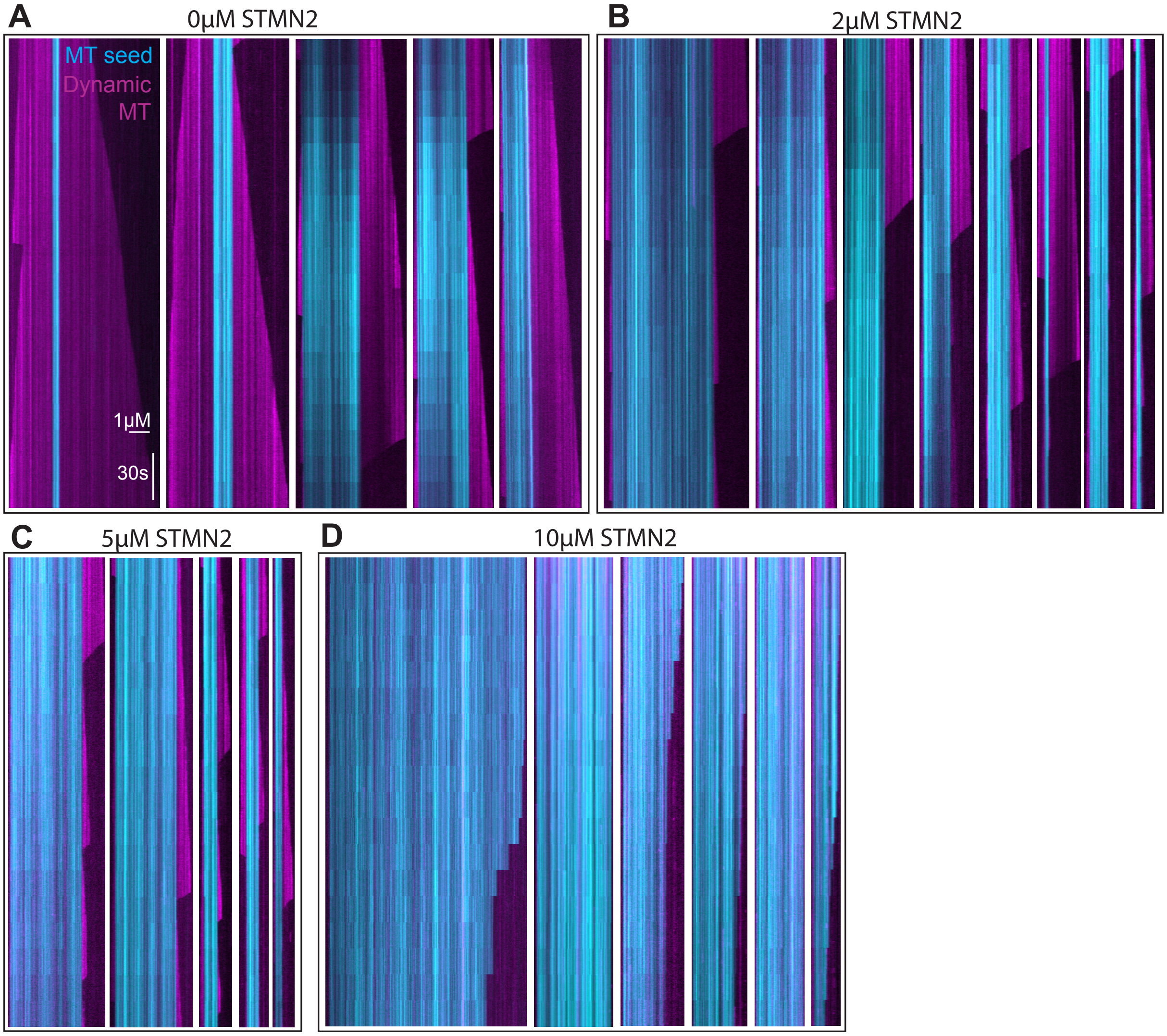

**Supplemental Figure 2. STMN2 is transported into distal axons and is positioned to regulate distal cytoskeletal organization; STMN2 depletion impacts tubulin and axon tips.**

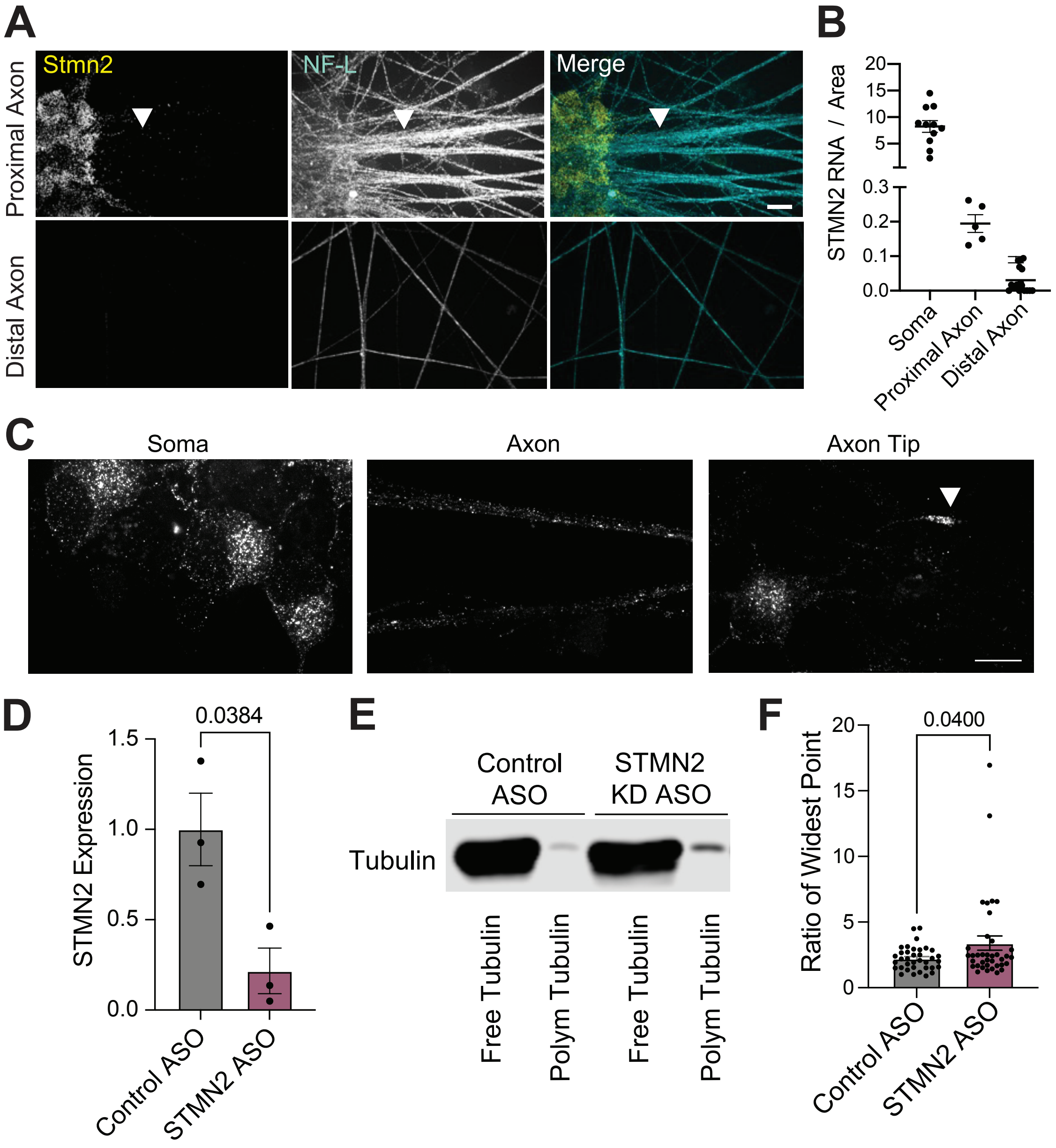

**Supplemental Figure 3. STMN2 overexpression alters microtubule polymerization.**

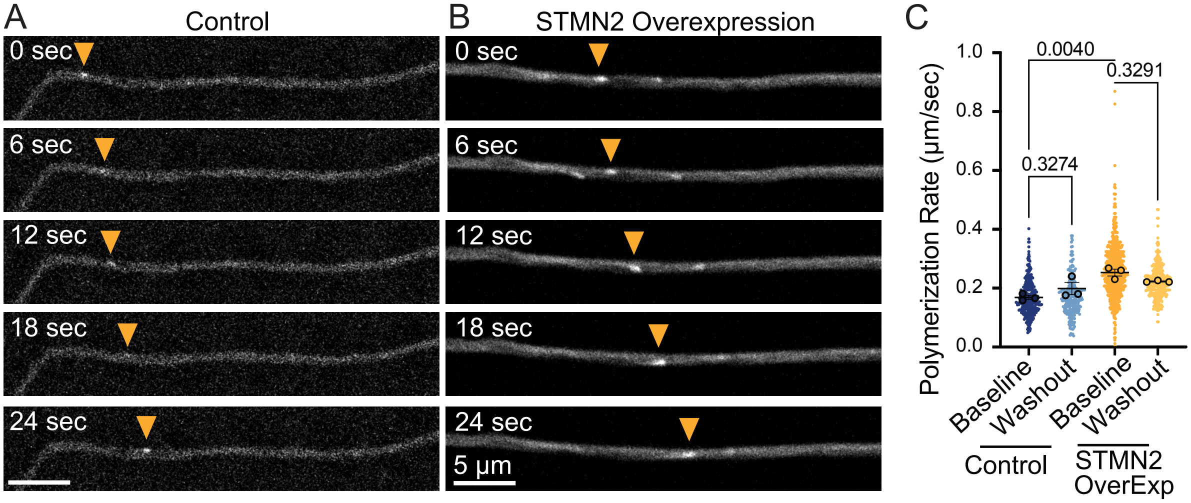


**Supplemental Figure 4. Mitochondrial localization with STMN2 knockdown.**

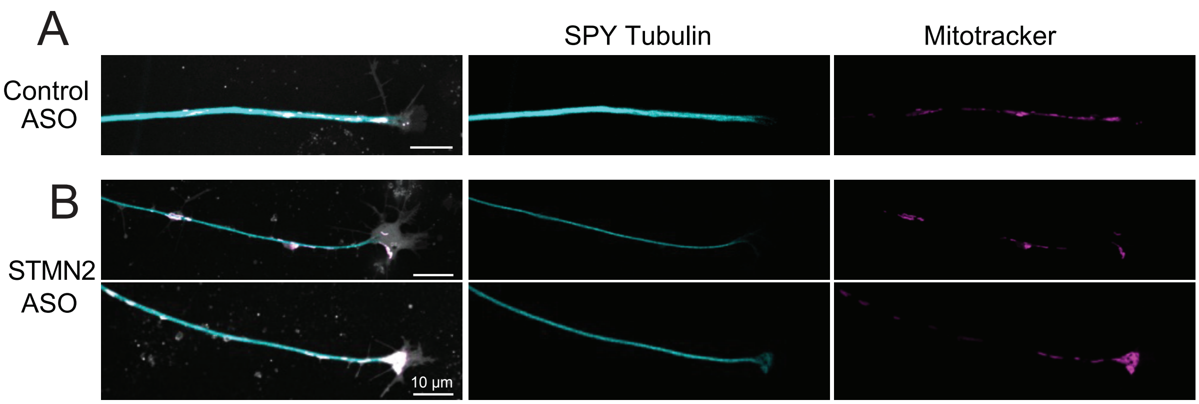
